# All-optical mapping of Ca^2+^ transport and homeostasis in dendrites

**DOI:** 10.1101/2024.07.16.603652

**Authors:** Rebecca Frank Hayward, Adam E. Cohen

## Abstract

Calcium mediates many important signals in dendrites. However, the basic transport properties of calcium in dendrites have been difficult to measure: how far and how fast does a local influx of calcium propagate? We developed an all-optical system for simultaneous targeted Ca^2+^ import and concentration mapping. We co-expressed a blue light-activated calcium selective channelrhodopsin, CapChR2, with a far-red calcium sensor, FR-GECO1c, in cultured rat hippocampal neurons, and used patterned optogenetic stimulation to introduce calcium into cells with user-defined patterns of space and time. We determined a mean steady-state length constant for Ca^2+^ transport *ϕ* ∼ 5.8 μm, a half-life for return to baseline *t*_1/2_ ∼ 1.7 s, and an effective diffusion coefficient *D* ∼ 20 μm^2^/s, though there were substantial differences in Ca^2+^ dynamics between proximal and distal dendrites. At high Ca^2+^ concentration, distal dendrites showed nonlinear activation of Ca^2+^ efflux, which we pharmacologically ascribed to the NCX1 antiporter. Genetically encoded tools for all-optical study of Ca^2+^ transport and handling provide a powerful capability for studying this important messenger.

## Introduction

Calcium is an important signaling molecule in all kingdoms of life.^1^ In mammals, Ca^2+^ is involved in many crucial functions, such as muscle contraction,^2^ hormone secretion,^3^ and neural activity and plasticity.^4^ The cytoplasmic Ca^2+^ concentration is determined by a complex interplay of transport across the plasma membrane, transport to and from internal stores in the endoplasmic reticulum (ER) and mitochondria, and buffering by Ca^2+^ binding proteins.

Ca^2+^ signaling is often compartmentalized to sub-cellular locations to achieve region-specific effects.^5–7^ This compartmentalization can occur over nanodomains associated with individual Ca^2+^ channels,^8^ microdomains associated with clusters of Ca^2+^ channels,^9^ or over larger regions defined by the geometry of the cell membrane, such as within presynaptic boutons,^10^ dendritic spines,^11^ primary cilia,^12^ or individual dendrite branches.^13,14^

To study sub-cellular Ca^2+^ transport and homeostasis, one would like to induce Ca^2+^ influx in arbitrary patterns of space and time, and to map the resulting time-dependent Ca^2+^ concentration profiles. A genetically encoded system would also permit targeting specific cell types. All-optical genetically encoded tools for targeted perturbation and measurement are a powerful approach to mapping spatiotemporal responses of cellular signals. For example, we have developed all-optical systems for perturbation and measurement of voltage,^15–17^ of cAMP,^18^ and of the embryonic morphogen Nodal.^19^

Here we co-expressed a blue-excited calcium-selective channelrhodopsin CapChR2^20^ with a farred fluorescent Ca^2+^ indicator protein FR-GECO1c^21^ in cultured rat hippocampal neurons. We used optical patterning to introduce local calcium fluxes and determined the resulting steady state length-constant for Ca^2+^ spread. We then used wide-area channelrhodopsin activation to map cell-wide Ca^2+^ recovery kinetics. We combined these measurements to infer the effective diffusion coefficient of Ca^2+^ ions in the dendrites. We then explored how blocking the sodium-calcium exchanger NCX1 affected Ca^2+^ transport and recovery after a perturbation.

### Simple model: 1-D reaction diffusion

Reaction-diffusion dynamics provide a conceptual framework for interpreting the experiments below, so we introduce the relevant formulas and terms here. Similar ideas have been presented elsewhere, e.g. Ref. ^22^. We treat each dendrite as a 1-D tube (we ignore branching). The Ca^2+^ dynamics then follow:

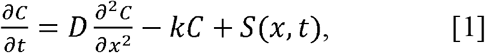

where *x* is the contour coordinate along the tube, *C* is the deviation from equilibrium Ca^2+^ concentration, *D* is an effective diffusion coefficient, and *k* is a rate constant for return of Ca^2+^ to equilibrium. *S*(*x,t*) is the source density, i.e. the time-dependent injection or removal of Ca^2+^ into the dendrite, such as by CapChR2. Here *D* can (and does) differ substantially from the free-solution diffusion coefficient, because it includes the effects of interactions with buffers, uptake and release from internal stores, occlusion by organelles, and transient trapping of Ca^2+^ ions in dendritic spines. Here *k* captures the effects of all Ca^2+^ transport processes which tend to return the Ca^2+^ concentration to its equilibrium value. This includes transport across the plasma membrane, and also uptake by internal stores.

Eq. 1 is a linear partial differential equation, which can be solved by standard techniques. A sum of two solutions is also a solution. It is therefore convenient to introduce the impulse response, or Green’s function, *G*(*x,t*). To calculate the concentration profile for an arbitrary time-dependent input *S*(*x,t*), one simply convolves *G*(*x,t*) with *S*(*x,t*). Under steady-state conditions, i.e. *S*(*x,t*) = *S*(*x*), the steady state Green’s function, *G*_*ss*_(*x*), is particularly simple:

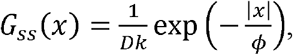

where the length constant 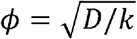. This Green’s function gives the steady state calcium concentration profile around a constant Ca^2+^ flux localized around *x* = 0, i.e. for *S*(*x*)= *δ*(*x*).

For a spatially homogeneous initial deviation in concentration *C*_0_, the time-dependent deviation in concentration is *C*(*t*) = *C*_0_exp (−*k t*). Thus by measuring length constant *ϕ* and recovery rate constant *k*, one can determine the effective diffusion coefficient. *D* =ϕ^2^*k* For a more detailed discussion of Eq. 1 and its time-dependent solutions, see e.g. Ref. ^18^.

The above framework is an oversimplification of reality. It assumes that the underlying dynamics are linear, spatially homogeneous, and constant in time. There are many reasons to doubt these assumptions. As Ca^2+^ concentration increases, intracellular buffers become saturated and the fraction of bound vs. free Ca^2+^ changes, leading to changes in *D*,^23^ a situation handled by nonlinear models of buffered diffusion.^24^ Ca^2+^ dynamics can also be nonlinear because of Ca^2+^-induced Ca^2+^ release or uptake by the endoplasmic reticulum^25,26^ and mitochondria^27^. Finally, the inhomogeneous distribution of mitochondria can cause localized anomalies in Ca^2+^ concentrations,^14^ the approximation of the dendrite as a featureless 1-D tube is clearly incorrect, and Ca^2+^ buffering and transport are subject to plasticity at multiple timescales.^28,29^ Despite being an oversimplification, the model above is useful for predicting scaling properties of Ca^2+^ transport and for identifying deviations from the model assumptions.

### Experiments

#### All-optical Ca^2+^ perturbation and measurement with genetically encoded tools

We created an all-optical system for mapping Ca^2+^ signaling by co-expressing an actuator and reporter of Ca^2+^ (Figure 1A). To introduce controlled perturbations to Ca^2+^, we used CapChR2-eGFP^20^, a Ca^2+^ selective cation channel with an activation peak at 475 nm. We first performed a detailed characterization of CapChR2 in HEK293T cells using whole-cell patch clamp electrophysiology (Figure 1B–F). CapChR2 was highly sensitive to blue light (intensity for 50% activation, I_50_ = 0.035 mW/mm^2^, Figure 1C). The opening and closing time constants at 37 °C and 1.45 mW/mm^2^ were τ_on_ = 19 ± 3 ms and τ_off_ = 81 ± 14 ms, (mean ± s.e.m. n = 3 cells). The kinetics were slower at lower intensity or lower temperature (Fig. 1D). CapChR2 had a positive reversal potential (10 mV, Figure 1E), consistent with preferential Ca^2+^ selectivity. CapChR2 showed negligible sag in photocurrent during sustained illumination (Figure 1F).

**Figure 1:**
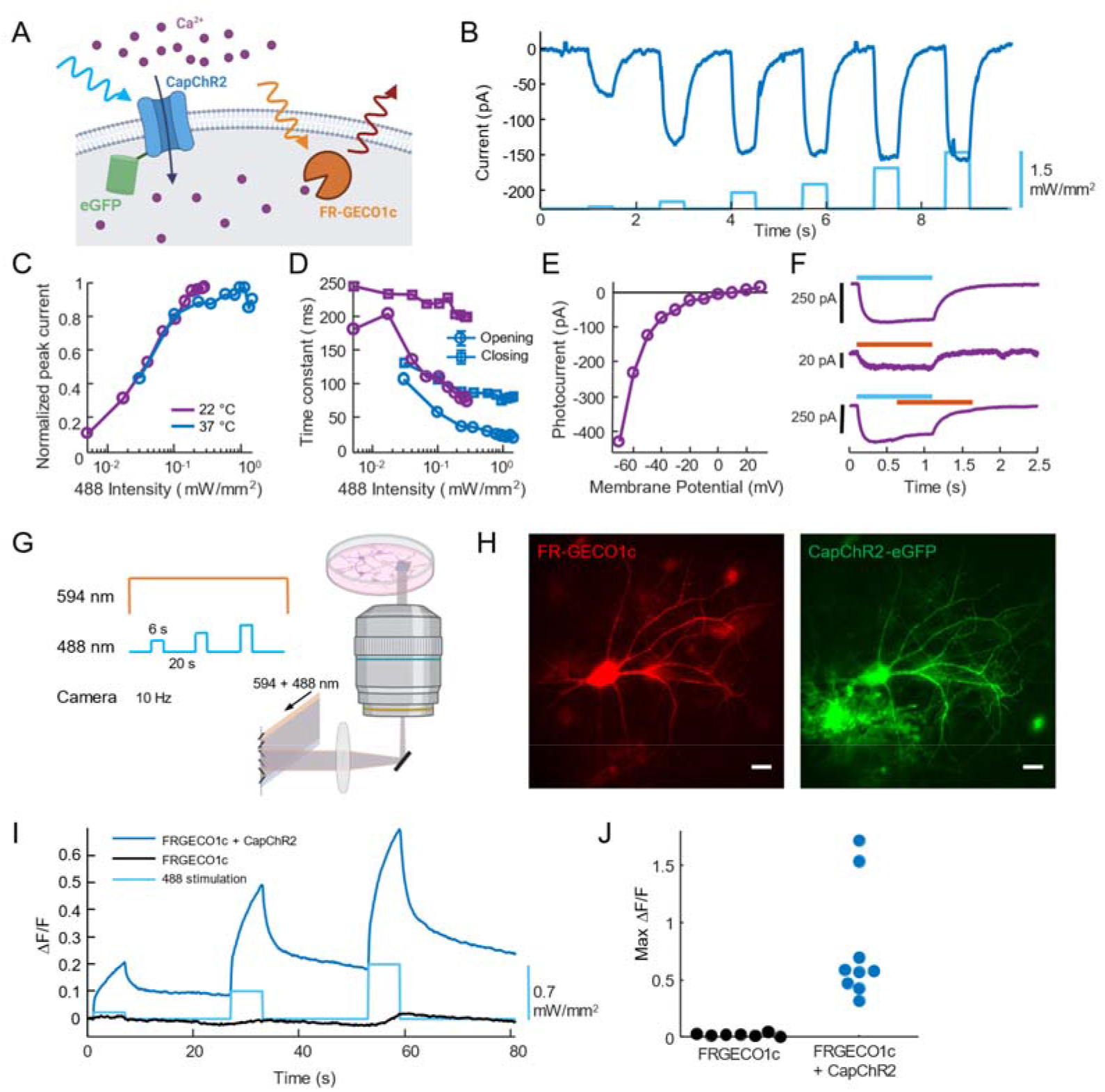
All-optical perturbation and measurement of Ca^2+^ dynamics. A. A blue light-activated Ca^2+^-selective Channelrhodopsin, CapChR2 is co-expressed with an orange-light excited Ca^2+^ sensor, FR-GECO1c. B. Example recording of CapChR2 photocurrents in a HEK293T cell at 37 ºC. Holding voltage -70 mV. C. Normalized peak photocurrent as a function of 488 nm illumination intensity. Blue: average of n = 3 cells at 37 ºC; purple: average of n = 2 cells at 22 ºC. D. CapChR2 opening and closing time constants as a function of 488 nm illumination intensity. Opening (circles) and closing (squares). Blue: average of n = 3 cells at 37 ºC; purple: average of n = 2 cells at 22 ºC. E. Photocurrent for CapChR2 in HEK293T cell as a function of holding voltage. Reversal potential was V_rev_ = 15 mV. Experiment at 22 ºC. F. Effect of 594 nm illumination on CaPChR2 gating. Illumination at 594 nm, mW/mm^2^ induced 8% photocurrent compared to illumination at 488 nm, mW/mm^2^. Co-illumination with 594 and 488 nm reduced photocurrent by 30 pA, or 9.6% relative to 488 nm alone. Experiment at 22 ºC. G. Protocol for simultaneous optogenetic activation of CaPChR2 and measurement of Ca^2+^ dynamics via fluorescence of FR-GECO1c. H. Example neuron showing expression of FR-GECO1c (red, left) and CapChR2-eGFP (green, right). Scale bars 20 μm. An astrocyte also expresses CapChR2-eGFP on the lower left. I. Representative fluorescence responses of FR-GECO1c at the soma in neurons with and without expression of CapChR2. Experiment at 37 ºC. J. Maximum fluorescence responses of FR-GECO1c at soma in neurons with (n = 9 neurons) and without (n = 7 neurons) CapChR2 expression. Experiment at 37 ºC.

To test whether the light used for imaging the red-shifted Ca^2+^ reporter affected the opsin, we compared CapChR2 photocurrents under blue (488 nm), orange (594 nm) and combined blue + orange illumination. At matched intensities of 0.22 mW/mm^2^, the photocurrent from orange light was only 8% of the photocurrent from blue light (Figure 1F). In some opsins, red or orange light can drive a transition from the open to the closed state.^30,31^ To test for this effect, we applied blue light (488 nm, 0.22 mW/mm^2^), and then after a 3.5 s delay, we applied orange light (594 nm, 0.21 mW/mm^2^) while maintaining the blue illumination (Figure 1F). The orange light induced a decrease in photocurrent of ∼10%. Since the orange illumination was constant and homogeneous in our measurements, this slight decrease in overall CapChR2 photocurrent was not consequential.

To map the CapChR2-evoked Ca^2+^ concentration profile, we used FR-GECO1c,^21^ a far-red Ca^2+^ reporter with an excitation peak at 596 nm. This reporter has previously been characterized in detail. The reported response properties relevant to our study are: calcium affinity 83 nM, maximum response amplitude (ΔF/F) 18, Hill coefficient 2, rise half-time 13 ms, decay half-time 156 ms.^21^

We then tested the combined system in cultured neurons. Cultured neurons were transduced with separate lentiviral vectors for FR-GECO1c and CapChR2-eGFP. We used a custom microscope (Methods, Figure 1G) in which 594 nm light and 488 nm light were combined, passed through an acoust-optical tunable filter, and patterned via a digital micromirror device (DMD, Figure 1G) which was re-imaged onto the sample. By rapidly alternating between orange and blue illumination and synchronously switching between two DMD patterns, we could create arbitrary and independent spatial patterns of orange and blue illumination. Provided that the alternation was faster than the biological response times of the sample, the flicker had negligible effect.

All experiments were performed in 2 μM TTX to block spiking, so that changes in intracellular Ca^2+^ were primarily mediated by CapChR2 currents. Two-color imaging under wide-field 488 nm and 594 nm illumination identified cells which co-expressed the actuator and reporter (Figure 1H). These cells showed a large increase of the FR-GECO1c signal in the soma during wide-area blue light stimulation, (ΔF/F = 0.77 ± 0.50, mean ± s.d., n = 9 neurons). In neurons expressing only FR-GECO1c the fluorescence change was much smaller (ΔF/F = 0.026 ± 0.014, mean ± s.d., n = 7 neurons, Figure 1I, J). These experiments established that optical stimulation of CapChR2 caused Ca^2+^ influx to the cytosol and a corresponding increase in FR-GECO1c fluorescence, while blue light stimulation evoked negligible optical crosstalk in FR-GECO1c alone.

#### Targeted stimulation probes Ca^2+^ length constants

We then used targeted optogenetic stimulation of small segments of dendrites to probe Ca^2+^ transport (Figure 2A), again in the presence of 2 μM TTX. We used synchronous alternation of the illumination wavelength and the DMD patterns at 10 Hz to target the blue light to small dendritic patches and to provide wide-area orange illumination (Figure 2B). The camera was also triggered synchronously with the illumination patterns (Figure 2C). The 50 ms interval between blue stimulation epochs was shorter than the measured off-time of CapChR2, so this protocol gave a quasi-continuous photocurrent and near-simultaneous wide-field Ca^2+^ imaging at an effective rate of 10 Hz and 50% duty cycles for imaging and stimulation. The protocol also included some frames of orange-only illumination at the beginning, to provide a fluorescence baseline, and some frames of blue-only illumination at the end, to measure blue-excited FR-GECO1c fluorescence for post-measurement crosstalk correction.

**Figure 2:**
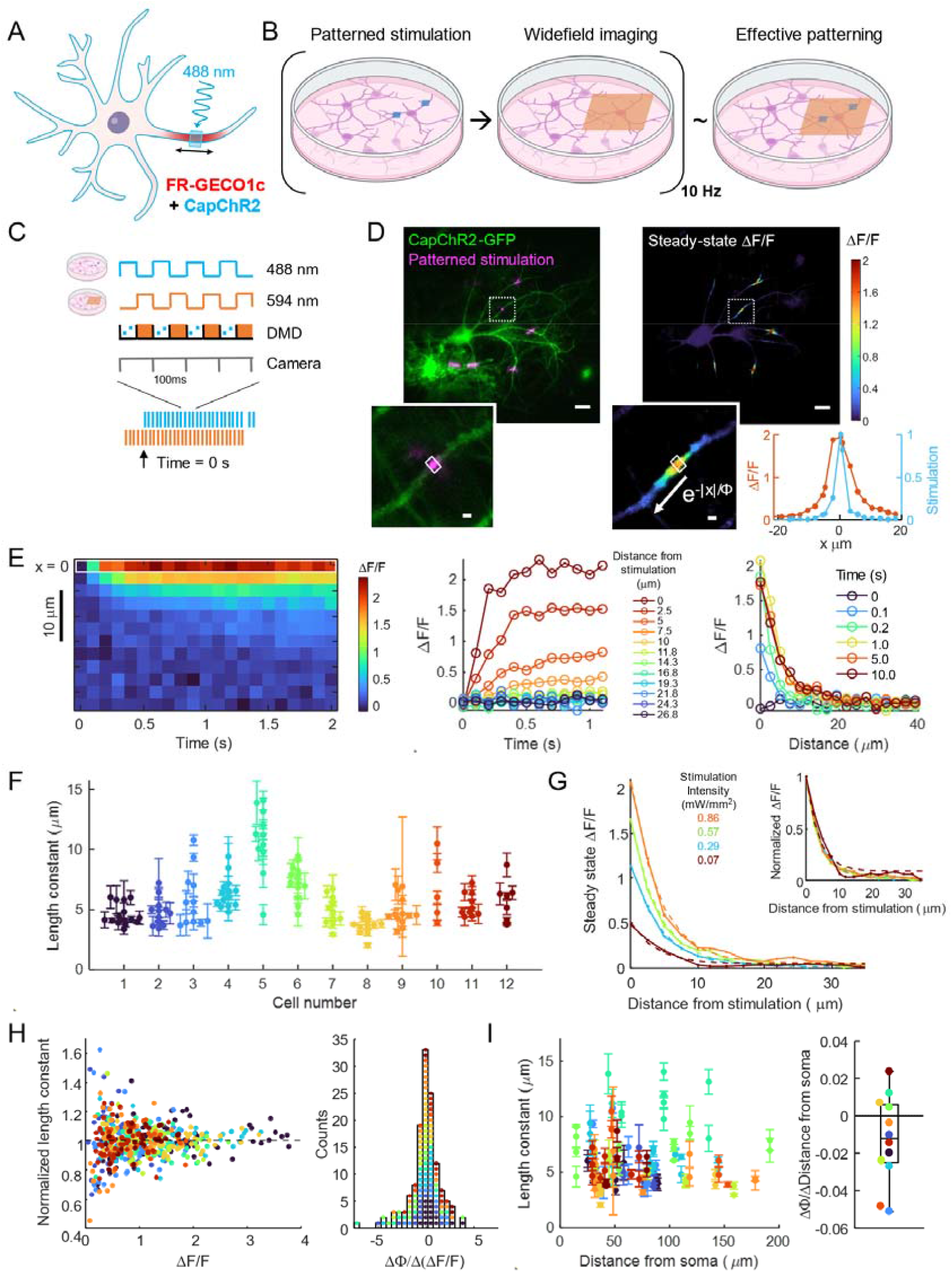
Measuring the Ca^2+^ length constant in dendrites. A. Experimental scheme to test the spread of Ca^2+^ around a local optical stimulation of CapChR2. B. Rapid (10 Hz) alternation of targeted blue stimulation and wide-field orange illumination gave an effective pattern consisting of both. C. Experiment timing. Camera triggers were synchronized to acquire 1 frame per illumination cycle. 10 s of orange-only imaging at the start of the experiment provided a measure of baseline fluorescence. Blue light-evoked autofluorescence was measured at the end of the experiment. D. Top left: example neuron showing GFP fluorescence under wide-field illumination (indicative of CapChR2, green), and GFP fluorescence under patterned optogenetic stimulation (indicative of stimulus pattern, magenta). Top right: steady-state ΔF/F of FR-GECO1c. Insets show stimulated regions. Bottom right: profile of stimulation (eGFP fluorescence, blue) and steady-state ΔF/F of FR-GECO1c (orange). Top scale bars 20 μm, bottom scale bars 2 μm. E. Example response along a dendrite to localized stimulation. Left: Kymograph. Center: ΔF/F vs. time at different distances. Right: ΔF/F vs. position at different times. F. Length constants for multiple stimulation spots and multiple cells. Error bars show standard deviation of n = 3-5 measurements on same stimulation spot. G. Steady-state ΔF/F vs. distance from stimulation for experiments with varying stimulus intensities. Solid lines: data; dotted lines: exponential fits. Inset: plots normalized to their maximum ΔF/F. H. Left: length constant normalized by the average over multiple stimulation intensities at the same stimulation spot, colored by cell (n = 12 neurons, n = 135 stimulation spots). Peak ΔF/F was used as a proxy for local stimulation strength. For each stimulation spot, a line was fitted to the normalized length constant vs. peak ΔF/F. Right: histogram of slopes for these fits. I. Left: length constant at each stimulation spot vs. the contour distance from the soma, colored by cell. For each cell the data were fit to a line. Right: slopes of change in length constant vs. distance from soma.

For each cell, we selected 3 – 6 well-separated dendritic regions to stimulate simultaneously. These regions were far enough apart that their calcium responses did not overlap. The FR-GECO1c fluorescence initially rose at the stimulus locations, and then spread, reaching a steady state after ∼1 s (Figure 2D,E, Supplementary Movies 1, 2). The steady-state Ca^2+^ distribution extended substantially beyond the stimulated zone, showing the diffusion of Ca^2+^ ions from the stimulation spot (Figure 2D). We fit the shape of the calcium concentration profile with an exponential decay, with a length constant *ϕ*. In control experiments with neurons expressing only FR-GECO1c, targeted blue stimulation did not evoke a detectable change in FR-GECO1c fluorescence (Fig. S1).

We measured the length constants from 135 local stimulation spots over 12 neurons for multiple stimulation intensities (Figure 2F). There was more cell-to-cell variability in *ϕ* between neurons than between spots within the same neuron. We speculate that this difference may be due to cell-to-cell differences in expression of Ca^2+^-transport proteins, leading to differences in efflux rate. The population-average length constant was 5.8 ± 2.3 μm (mean ± s.d.), with cell-average length constants ranging from 3.7 ± 0.7 μm to 10.8 ± 2.7 μm. This result is broadly consistent with the simulation results of Biess and coworkers, who predicted a steady-state full-width at half-maximum (FWHM) of the Ca^2+^ signal of 5 μm, corresponding to an exponential decay length constant of 3.6 μm.^32^

We then varied the stimulation intensity from 0.07 to 0.86 mW/mm^2^ to test for concentration dependence of Ca^2+^ transport (Figure 2G). Steady-state Ca^2+^ concentration increased with stimulation intensity, as expected, but when the spatial profiles were normalized to their respective peaks, they appeared to have similar length constant (Figure 2G inset). Comparisons between stimulus locations were complicated by the large cell-to-cell variations in *ϕ*, and also by the fact that there were uncontrolled variations in optogenetic stimulation strength between regions. This spot-to-spot variability came from variations in CapChR2 expression level and from variations in the membrane area underlying each stimulation spot.

We thus focused on within-spot relations between Ca^2+^ level and length constant. For each spot, we calculated a *ϕ*_norm_= *ϕ*/⟨*ϕ*⟩ where ⟨*ϕ*⟩ is the average across stimulation intensities of *ϕ* for that spot. We then plotted *ϕ*_norm_ vs. ΔF/F (Figure 2H). We did not observe a global trend. We then fit the slope of *ϕ*_norm_ vs. ΔF/F for each stimulation spot, and plotted a histogram of the slopes. This histogram was centered at zero. Together, these results establish that we did not detect a relationship between the amplitude of the local Ca^2+^ transient and the steady-state length constant.

We also asked whether the length constant depended on contour distance, *d*, from the soma (Figure 2I). To account for the cell-to-cell variability in *ϕ*, we separately fit a line to *ϕ* vs. *d* for each cell. Length constants were slightly shorter in more distal dendrites, with a median slope of -0.012 (dimensionless; quartiles = [-0.025, 0.0058]). For example, at a slope of -.01, the length constant would be 1 μm shorter at 100 μm from the soma than near the soma. This distance dependence is consistent with a membrane-based Ca^2+^ efflux mechanism: smaller-diameter dendrites have higher surface-to-volume ratio, leading to more rapid efflux and hence shorter length constants. Our measurements are qualitatively consistent with the results of Korkotian and Segal, who measured that under impulsive Ca^2+^ uncaging the half-decay lengths were 1.5 μm in “thick” and 1.2 μm in “thin” dendrites (definitions not specified).^33^

#### Wide-field stimulation probes Ca^2+^ dynamics

We used wide-field CapChR2 stimulation and wide-field FR-GECO1c imaging to map the cell-wide Ca^2+^ response kinetics to perturbations (Figure 3A). As before, we used TTX (2 μM) to suppress spiking. The stimulus waveform comprised three pulses of 6 s duration, with intensity increasing from 0.07 – 0.57 mW/mm^2^, and with 20 s rest between pulses. At the end of the experiment, we turned off the orange light and recorded autofluorescence excited by the blue light alone for each stimulation intensity. These crosstalk signals were then subtracted from the data acquired with the orange laser on.

**Figure 3:**
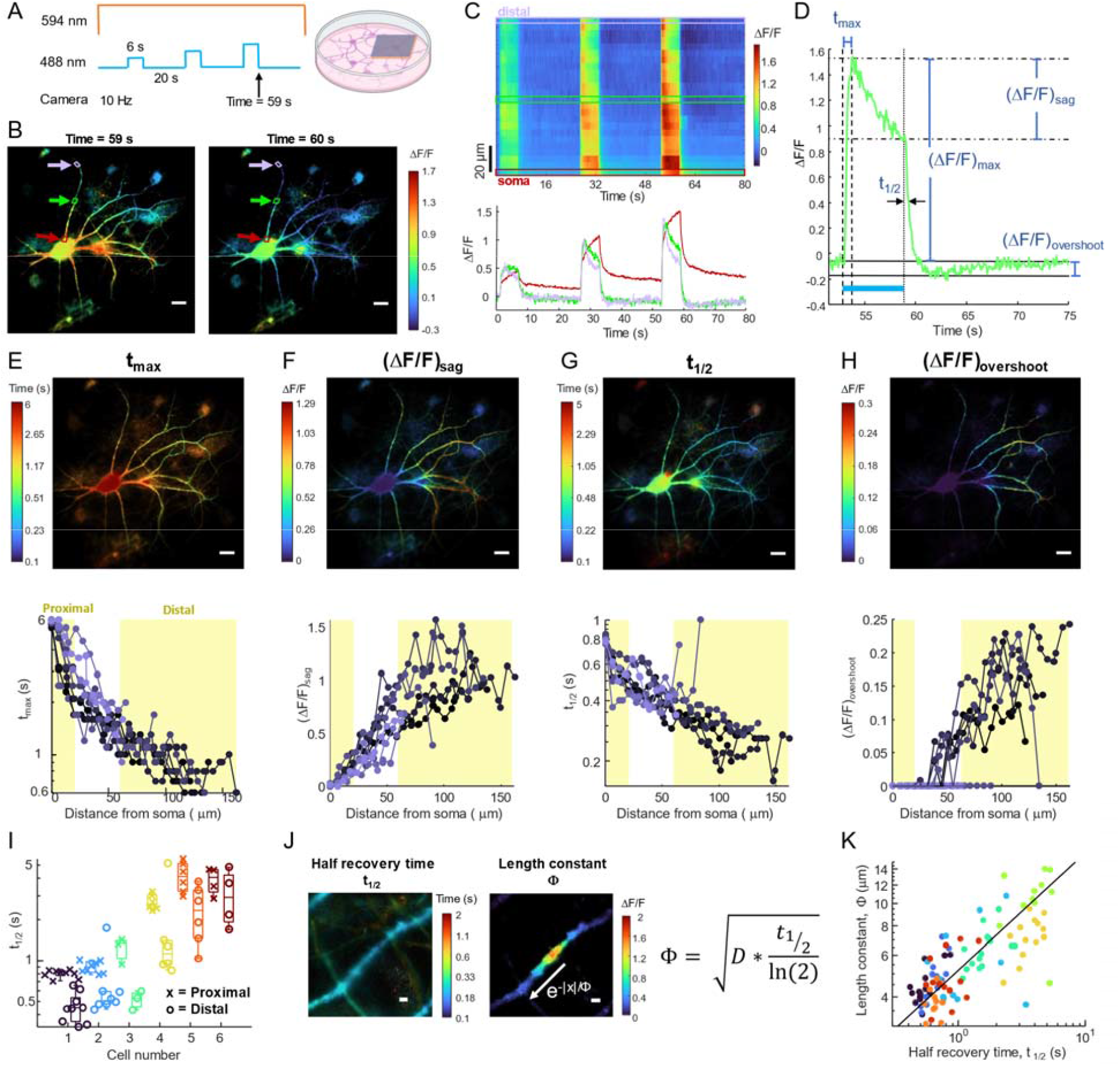
Ca^2+^ dynamics under wide-area stimulation. A. Wide-area optogenetic stimulation induces global elevation in Ca^2+^. B. Map of FR-GECO1c fluorescence. Left: at the end of the strongest stimulus pulse. Right: 1 s after the end of the stimulus. Scale bars 20 μm. C. Top: kymograph of ΔF/F (top) along dendrite shown by arrows in B. Bottom: fluorescence at locations marked by arrows in B. D. Example plot of ΔF/F showing the definitions of t_max_, (ΔF/F)_sag_, t_1/2_, and (ΔF/F)_overshoot_. E-H. Top: maps of the indicated kinetic parameters for a representative cell. Bottom: plotted as a function of contour distance from the soma. I.Indicated kinetic parameters averaged over proximal and distal dendrites across multiple cells. J.Left: map of the half recovery time, and right: the steady state ΔF/F in an example stimulation spot. Right: Predicted relation between recovery time, length constant, and diffusion coefficient. Scale bars 2 μm. K.Length constant, ϕ, vs. half-recovery time *t*_1/2_. Median diffusion coefficient is shown as black line.

The cytosolic Ca^2+^ grew during each blue stimulus and returned toward baseline after, but the dynamics showed striking variations with distance from the soma (Figure 3B, C, Supplementary Movie 3). To quantify these dynamics, we defined several parameters, shown in Fig. 3D. Figures 3E-H show maps of the Ca^2+^ dynamics for a representative neuron, and quantification of these dynamics as a function of distance from the soma. Upon stimulus onset, the fluorescence in distal branches peaked in t_max_ < 1 s and then started to decrease, whereas the fluorescence in the soma and nearby dendrites continued to grow for the full 6 s stimulus. The amount of sag in the fluorescence after the peak, (ΔF/F)_sag_, was greater in the distal branches than in and near the soma (where it was zero). Upon termination of the stimulus, the half-recovery time of the Ca^2+^ signal was faster in the distal dendrites (*t*_1/2_ ∼ 0.3 s) than near the soma (*t*_1/2_ ∼ 0.6 s). In the distal branches, the FR-GECO1c fluorescence then decreased below its initial basal value, before increasing back to baseline, while there was no fluorescence overshoot during recovery in and near the soma.

Figure 3I shows a summary of the proximal and distal recovery times across neurons and Fig. S2 shows the distributions of the other kinetic parameters across cells. We defined the proximal dendrites to have *d* < 20 μm from the soma, and distal dendrites as *d* > 60 μm from the soma. As with the length constants, there was substantial cell-to-cell variability in the Ca^2+^ recovery kinetics, although the proximal-to-distal trends in the dynamics were consistent.

We propose that the proximal-to-distal variation in Ca^2+^ dynamics can be explained by the greater surface-to-volume ratio in the more distal dendrites^34,35^. For a given current density (influx or efflux) across the membrane, the rate of change of concentration in the dendrite scales as 1/*r*, where *r* is the radius of the dendrite. This scaling accounts for the shorter time to peak, *t*_max_, in the smaller diameter distal dendrites. This scaling also likely accounts for the faster recovery time, *t*_1/2_, in the more distal dendrites, where the efflux is through the Ca^2+^ ATP-ase (PMCA) and Na^+^-Ca^2+^ Exchanger (NCX)^36^.

The sag during the stimulus epoch and the post-stimulus overshoot cannot be explained within the linear reaction-diffusion model of Eq. 1. Since the sag and the overshoot colocalized within cells and co-varied in magnitude between cells, we propose that these are likely through a similar mechanism, e.g. a Ca^2+^ efflux mechanism that is activated at high Ca^2+^ concentration and remains active for a time after the Ca^2+^ concentration returns to baseline. We return to this mechanism in the next section.

The measurements of length constants (Fig. 2) and recovery kinetics (Fig. 3) were performed on the same cells, so we combined these measurements to estimate an effective diffusion coefficient, *D*. For each location where we had applied targeted stimulation and measured a length constant, we also determined the Ca^2+^ recovery time constant *t*_1/2_ from the wide-field stimulation data. In a simple model of first-order efflux, the recovery rate constant is *k* = ln(2)/*t*_1/2_. While there was substantial cell-to-cell variation in *ϕ* (e.g. Fig. 2D,F) and in *t*_1/2_ (Fig. 3G,I), on a log-log plot of *ϕ* vs. *t*_1/2_, the data clustered along a line. The inferred effective diffusion coefficient was *D* =19.0 μm^2^/s (95 % Bootstrap Confidence Interval [16.9, 21.4], n = 8 neurons, 102 stimulation spots; Figure 3J,K).

#### NCX1 mediates nonlinear recovery kinetics

The sodium-calcium exchanger NCX1 is expressed in hippocampal pyramidal cell dendrites *in vivo*^37^ and in culture.^38^ NCX1 is a low-affinity (0.2 – 10 μM),^39^ high-capacity electrogenic Ca^2+^ transporter, whose primary role is to mediate rapid efflux of Ca^2+^ following an action potential.^40^

The activity of this antiporter is dynamically regulated by Ca^2+^ binding to allosteric regulatory sites, as well as by many other intracellular signals.^41^ We thus hypothesized that the nonlinear dynamics in distal dendrites (sag during stimulation and overshoot following stimulation) might be attributable to Ca^2+^-mediated allosteric enhancement of NCX1 activity.

We compared optogenetically evoked Ca^2+^ dynamics before and after blocking NCX1 with benzamil (30 μM; Fig. 4A)^42^. In both proximal and distal dendrites, benzamil increased the time to maximum fluorescence, (*t*_max_: 3.5 ± 1.7 s before, 4.1 ± 1.5 s after, P = 6.8e-8, paired two-sided t-test, *n* = 25 dendrites, 5 neurons), and increased the post-stimulation recovery time, (*t*_1/2_: 1.8 ± 1.4 s before, 2.0 after, P = 1.8e-9, paired two-sided t-test). In distal dendrites, benzamil also decreased the sag during stimulation, ((ΔF/F)_sag_: 0.43 ± 0.27 before, 0.26 ± 0.23 ± 1.4 s after, P = 1.1e-9, paired two-sided t-test, *n* = 25 dendrites, 5 neurons), and largely eliminated the overshoot before return to baseline ((ΔF/F)_overshoot_: 0.058 ± 0.028 before, 0.034 ± 0.029 after, P = 0.0065, n = 12 dendrites, 3 neurons which displayed overshoot behavior before drug addition, paired twosided t-test; Fig. 4B). Together these results support the hypothesis that NCX1 acts as a nonlinear efflux pump, activated at high Ca^2+^ and slowly inactivating upon return to baseline Ca^2+^.

**Figure 4:**
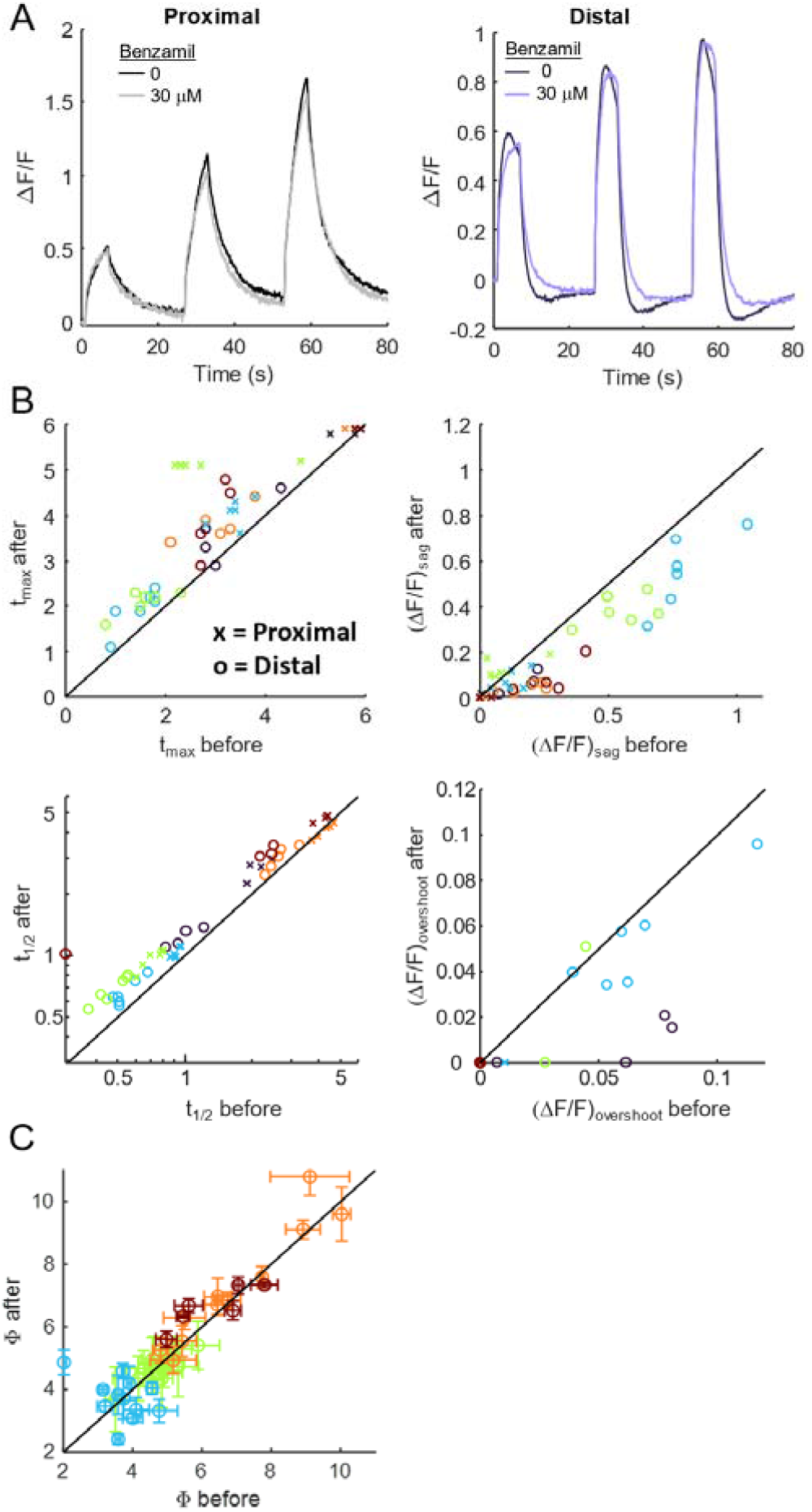
Effect of NCX block on calcium length constant and recovery kinetics. A. Left: ΔF/F of proximal and right: distal section of example dendrite before (dark line) and after (light line) addition of benzamil to block NCX1. B. Effect of benzamil on t_max_ (top left), (ΔF/F)_sag_ (top right), t_1/2_ (bottom left), and (ΔF/F)_overshoot_ (bottom right) in distal and proximal regions of dendrites (n = 25 dendrites, 5 neurons). C. Comparison of length constants (n = 41 dendrites, 4 neurons) before and after NCX1 block. Error bars show standard deviation over 3 measurements

We did not observe an effect of benzamil on the length constant in either proximal or distal dendrites (Fig. 4C). The measured increase in recovery time constant, *t*_1/2_, was ∼27%, implying an increase in length constant of ∼13%. This change was likely too small for us to detect.

## Discussion

We combined a blue-excited Ca^2+^-permeable channelrhodopsin with an orange-excited fluorescent Ca^2+^ indicator for all-optical mapping of Ca^2+^ response-functions in dendrites. We measured length constants *ϕ* ∼ 5.8 μm and recovery times *t*_1/2_ ∼ 1.7 s. From these, we calculated an effective Ca^2+^ diffusion coefficient, *D* ∼20 μm^2^/s. Though there was substantial branch-to-branch and cell-to-cell variation in *ϕ* and *t*_1/2_, *D* was comparatively constant.

Free Ca^2+^ in water has a diffusion coefficient of *D* = 778 μm^2^/s.^43^ Several prior studies reported measurements of Ca^2+^ diffusion in cellular contexts. In a classic experiment in *Xenopus* egg extract, Allbritton, Meyer and Streyer measured a diffusion coefficient of 13 μm^2^/s for a radioactive ^45^Ca^2+^ tracer at a basal concentration of 90 nM^23^, which grew to 65 μm^2^/s at a basal concentration of 1 μM. However, in the Allbritton experiments, Ca^2+^ uptake and release from mitochondria and ER were pharmacologically blocked, so the roles of these stores on effective transport was not determined. A further critique of this result was that, by not mapping the concentration of the non-radioactive basal Ca^2+^ ions, the measurements ignored the contribution of these ions as they were displaced from buffers by ^45^Ca^2+^ tracers, i.e. that “messages diffuse faster than the messenger”.^44^ Also, in the long-and-thin geometry of dendrites, membranous organelles might occlude Ca^2+^ transport,^32^ and dendritic spines might trap Ca^2+^ ions,^45^ effects missing from purified cytoplasm. Despite these caveats, our results are broadly consistent with the results of Allbritton and coworkers, interpolated to a basal Ca^2+^ concentration of 100 – 200 nM.

More recent measurements used fluorescent Ca^2+^ indicators to measure transport of all Ca^2+^ ions. Murthy and coworkers measured the spread of Ca^2+^ signals in response to quantal activation of dendritic spines in cultured hippocampal pyramidal cells, and determined an effective diffusion coefficient of 21 μm^2^/s (95% CI: 10 – 36 μm^2^/s),^22^ remarkably close to our result. Biess and coworkers used photochemical uncaging of an inert dye to determine that geometrical effects alone (i.e. occlusion by organelles, trapping in spines, increased cytoplasmic viscosity) led to a ∼20-fold decrease in *D* in hippocampal dendrites compared to free solution.^32^ Thus, one would expect that once buffering is taken into account *D*_Ca2+_ < 778/20 = 39 μm^2^/s in dendrites.

Noguchi and coworkers used two-photon glutamate uncaging and calcium imaging in rat CA1 pyramidal cells in acute hippocampal slices to estimate a length constant of 1.6 – 1.9 μm and a calcium diffusion coefficient of 12 μm^2^/s.^46^ Korkotian and Segal used photochemical uncaging of Ca^2+^ directly in hippocampal dendrites and determined that Ca^2+^ transients spread over 3 – 3.5 μm, but did not extract a diffusion coefficient. A limitation of the above studies is that because they relied on an impulsive introduction of Ca^2+^ into the dendrite (either by synaptic input or by uncaging), it was difficult to distinguish between Ca^2+^ efflux and diffusion-driven dilution. This limitation may explain why other studies reported a modestly shorter length constant than we observed.

Using an analogous experiment with a blue-light activated adenylyl cyclase and a red-shifted cAMP reporter, we previously measured a dendritic length constant of cAMP transport *ϕ*_cAMP_ ∼30 μm, and a dendritic diffusion coefficient of *D*_cAMP_ ∼ 120 μm^2^/s.^18^ Allbritton and coworkers measured diffusion of inositol 1,4,5-trisphosphate (IP_3_) in free cytosol and determined *D*_IP3_ ∼ 283 μm^2^/s. They estimated a range of action of *ϕ*_IP3_ ∼ 24 μm, though both numbers would be expected to be smaller in dendrites due to the presence of occlusions and traps. Using a blue-light activated channelrhodopsin and a red-shifted voltage indicator, we previously measured a dendritic length constant of voltage transport *ϕ*_V_ ∼255 μm.^17^ This hierarchy of signaling length-scales is an important aspect of modeling neural dynamics.

Ca^2+^ handling differs between neuronal sub-types,^45,47–49^, and is also fundamentally important in many non-neuronal cell types such as astrocytes^50^, cardiomyocytes,^51^ and immune cells.^52^ Ca^2+^ signaling is also disrupted in many diseases of the nervous, cardiac, and immune systems. The tools we introduce here are expected to be compatible with a wide diversity of cell types. The tools could find application in high-throughput screens such as are used in drug development, in detailed mechanistic studies in cultured cells, and in applications in tissue and *in vivo*.

## Acknowledgments

We thank Katherine Xiang, Andrew Preecha and Shahinoor Begum for helpful discussions and technical assistance. This work was supported by a Vannevar Bush Faculty Fellowship, NIH grants 1-R01-MH117042, and 1-R01-NS126043, and Chan Zuckerberg Initiative dynamic imaging grant 2023-321177.

## Materials and Methods

### HEK293T cell culture

HEK293T cell lines were maintained at 37□°C, 5% CO_2_ in Dulbecco’s Modified Eagle Medium (DMEM) supplemented with 10% fetal bovine serum, 1% GlutaMax-I, penicillin (100 U/mL), streptomycin (100 μg/mL). For maintaining or expanding the cell culture, we used TC-treated culture dishes (Corning). For all imaging experiments, cells were plated on poly-(D)-lysine (PDL)-coated glass-bottomed dishes (Cellvis, Cat.# D35-14-1.5-N).

### Neuron culture

Primary E18 rat hippocampal neurons (fresh, never frozen, BrainBits #SDEHP) were dissociated following vendor protocols and plated in PDL-coated glass bottom dishes (Cellvis, Cat.# D35-14-1.5-N) at 21k/cm^2^.

### Cloning

CAG-FR-GECO1c was a gift from Robert Campbell (Addgene plasmid # 163682) and CapChR2_pmScarlet-N1 was a gift from Peter Hegemann (Addgene plasmid # 188032). Each gene was cloned into a lentiviral backbone with CMV promoter and can be found on Addgene.

### Lentivirus preparation

All lentivirus preparations were made in house. HEK293T cells were co-transfected with the second-generation packaging plasmid psPAX2 (Addgene #12260), envelope plasmid VSV-G (Addgene #12259) and transfer plasmids at a ratio of 9:4:14. For small batches of virus, we used 2.7 μg total plasmids and ∼300k cells in 35-mm dish. Some viruses were concentrated 1:10 using Lenti-X Concentrator (Takara Cat. # 631232) following vendor protocols. Quantities of virus used are quoted using volumes prior to concentration.

### Expression of optogenetic actuators and reporter

For characterization of CapChR2 photocurrents, HEK293T cells were transduced at least 2 days before imaging with 50-200 μL of lentivirus encoding CapChR2-eGFP.

Neurons were transduced after 6-10 days in culture with 200 μL lentivirus encoding FR-GECO1c driven by the CMV promoter and 50-200 μL of CapChR2-eGFP, also driven by the CMV promoter. Functional imaging was performed after 12-20 days in culture.

### Sample preparation for imaging

Before optical stimulation and imaging, 35□mm dishes were washed with 1□mL PBS to remove residual culture medium, then filled with 2□mL extracellular (XC) buffer containing (in mM): 125 NaCl, 2.5 KCl, 2 CaCl_2_, 1 MgCl_2_, 15 HEPES, 25 glucose (pH 7.3). All imaging and electrophysiology were performed using this XC buffer. For all experiments in neurons, we added 10 μM NBQX and 25 μM AP-V to block synaptic transmission, and 2 μM TTX to block spiking. For drug-addition experiments the stage-top incubator was briefly opened after the first imaging run for addition of 30 μM Benzamil (Tocris 3380) before commencing the second imaging run.

### Electrophysiology

For patch clamp measurements, filamented glass micropipettes (WPI) were pulled to a resistance of 5–10□MΩ and filled with internal solution containing (in mM) 6 NaCl, 130 K-aspartate, 2 MgCl_2_, 5 CaCl_2_, 11 EGTA, and 10 HEPES (pH 7.2). The patch electrode was controlled with a low-noise patch clamp amplifier (Axon Instruments MultiClamp 700B). Current traces were collected in voltage clamp mode. The electrophysiology data were filtered with a moving average of 4 ms to reduce noise. The time constants were fit using single exponentials. In plots with multiple wavelengths of stimulation, the currents were normalized to the peak current for 488 nm stimulation.

### Combined optogenetic stimulation and imaging

Experiments were conducted on a home-built inverted fluorescence microscope equipped with 488 nm and 594 nm laser lines and a scientific complementary metal-oxide semiconductor (CMOS) camera (Hamamatsu ORCA-Flash 4.0).

Beams from both lasers were combined using dichroic mirrors and sent through an acousto-optic tunable filter (Gooch and Housego TF525-250-6-3-GH18A) for temporal modulation of intensity of each wavelength. The beams were then expanded and sent to a DMD (Vialux, V-7000 UV, 9515) for spatial modulation. The beams were focused onto the back-focal plane of a 60×/1.2-NA water-immersion objective (Olympus UIS2 UPlanSApo ×60/1.20W). Fluorescence emission was separated from laser excitation using a dichroic mirror (405/488/594). Imaging of FR-GECO1c fluorescence was performed with 594 nm laser at illumination intensity 0.21 mW/mm^2^. Stimulation of CapChR2 was performed with 488 nm laser at illumination intensities of 0.005 - mW/mm^2^. Stimulation spots had lengths of 2-10 μm. Temperature was maintained using a covered stage-top incubator (Tokai Hit INUBG2ATFP-WSKM).

### Data Analysis

All data were processed and analyzed in MATLAB. At the start of each measurement, prior to any optogenetic stimulation, the orange laser was turned on and the sample allowed to equilibrate for 5 s. Then baseline fluorescence, F_0_, was measured via the average fluorescence during 4.5 s of 594 nm-only illumination. To correct for blue light-induced autofluorescence in the FR-GECO1c channel, at the end of the experiment images were acquired with blue-only illumination (orange light off). These images were then subtracted from the corresponding frames acquired during simultaneous optogenetic stimulation and Ca^2+^ imaging.

Individual cells expressing the desired constructs were selected by hand and fluorescence waveforms were calculated by averaging pixels in the selected region and subtracting the background, which was computed by taking the 20^th^ percentile of fluorescence intensity for pixels in a dark region of the field of view during imaging.

To determine steady-state Ca^2+^ length constant, the mean fluorescence profile was measured starting at the edge of the stimulation spot, from 6 - 19 s after blue stimulus onset. The profile was fit using a single exponential + constant.

To characterize the Ca^2+^ responses to wide-area stimulation, t_max_ was calculated as the time from the start of stimulation until maximum ΔF/F. (ΔF/F)_sag_ was calculated as the difference between ΔF/F at t_max_ vs. at end of stimulation. *t*_1/2_ was calculated as the time where ΔF/F was halfway between ΔF/F at the end of the stimulation and the baseline value after 20 s of relaxation. Linear interpolation was used to improve the time resolution in estimating *t*_1/2_. (ΔF/F)_overshoot_ was calculated as the difference between the minimum of ΔF/F during the recovery phase and the ΔF/F baseline after smoothing with a gaussian-weighted average over 4 s windows. (ΔF/F)_overshoot_ was only accepted as genuine if it was larger than the standard deviation in the ΔF/F baseline. Proximal and Distal measurements were calculated by averaging the ΔF/F from all regions 0 – 20 μm from soma and beyond 60 μm from soma, respectively.

For measurements of Ca^2+^ dynamics before and after addition of benzamil, some acquisitions had small movements between the two sets of measurements. The raw data were aligned using image registration functions in Matlab to ensure the same spatial regions are used pre- and post-drug.

## Supplementary movie captions

**Movie S1**. Ca^2+^ response of a single dendritic branch to local optogenetic stimulation.

**Movie S2**. Ca^2+^ responses across two neurons to targeted optogenetic stimuli.

**Movie S3**. Ca^2+^ responses to wide-area optogenetic stimulation.

**Supplemental Figure 1:**
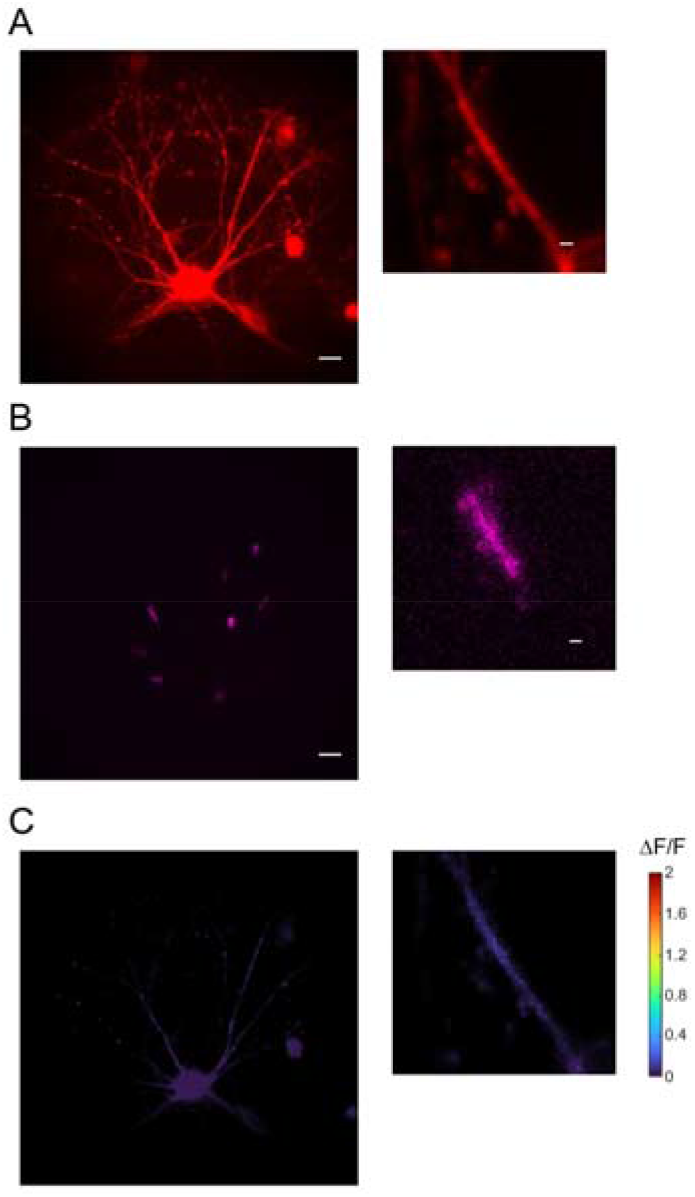
Effect of optical stimulation on neurons expressing sensor only. A. FR-GECO1c expression of example cell (left, scale bar 20 μm) with zoom in on example dendrite (right, scale bar 2 μm). B. Local stimulation spots on example cell (left) with zoom in on example dendrite (right). C. Steady-State ΔF/F of example cell (left) with zoom in on example dendrite (right).

**Supplemental Figure 2:**
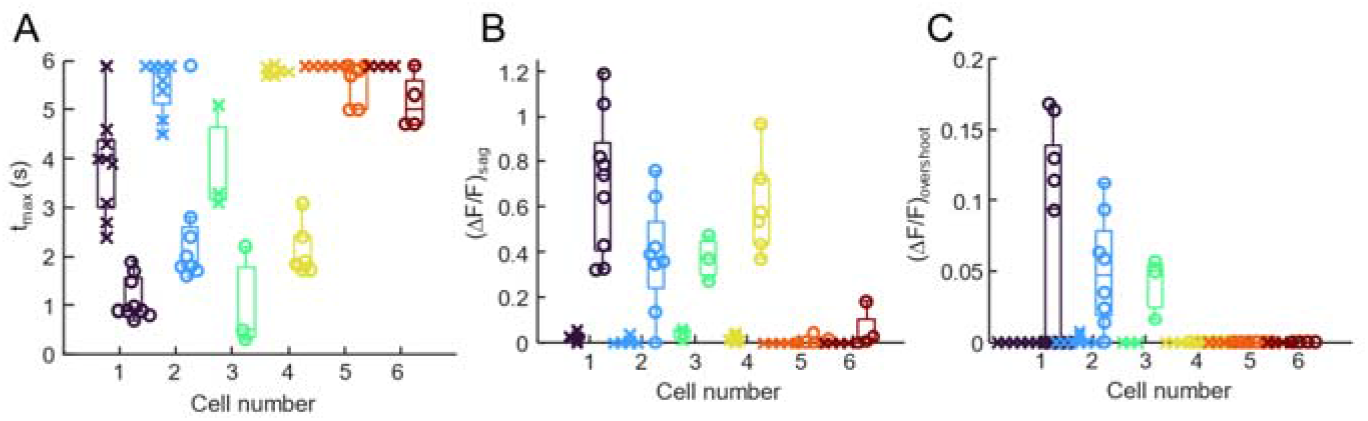
Cell-to-cell variability in parameters of Ca^2+^ dynamics. D. t_max_ in proximal and distal dendrites across multiple cells. E.(ΔF/F)_sag_ in proximal and distal dendrites across multiple cells. F.(ΔF/F)_overshoot_ in proximal and distal dendrites across multiple cells.

